# Electronic Mapping of a Bacterial Genome with Dual Solid-State Nanopores and Active Single-Molecule Control

**DOI:** 10.1101/2021.10.29.466509

**Authors:** Arthur Rand, Philip Zimny, Roland Nagel, Chaitra Telang, Justin Mollison, Aaron Bruns, Emily Leff, Walter Reisner, William B. Dunbar

## Abstract

We present the first electronic mapping of a bacterial genome using solid-state nanopore technology. A dual-nanopore architecture and active control logic are used to produce single-molecule data that enables estimation of distances between physical tags installed at sequence motifs within double-stranded DNA (dsDNA). Previously developed dual-pore “DNA flossing” control generates multiple scans of tagged regions of each captured DNA. The control logic was extended here in two ways: first, to automate “zooming out” on each molecule to progressively increase the number of tags scanned during DNA flossing; and second, to automate recapture of a molecule that exited flossing to enable interrogation of the same and/or different regions of the molecule. New analysis methods were developed to produce consensus alignments from each multi-scan event. The combined multi-scanning and multi-capture method was applied to the challenge of mapping from a heterogeneous mixture of single-molecule fragments that make up the *Escherichia coli* (*E. coli*) chromosome. Coverage of 3.1× across 2,355 resolvable sites (68% of reference sites) of the *E. coli* genome was achieved after 5.6 hours of recording time. The recapture method showed a 38% increase in the merged-event alignment length compared to single-scan alignments. The observed inter-tag resolution was 150 bp in engineered DNA molecules and 166 bp natively within fragments of *E. coli* DNA, with detection of 133 inter-site intervals shorter than 200 bp in the *E. coli* reference map. Proof of concept results on estimating distances in repetitive regions of the *E. coli* genome are also provided. With an appropriately designed array and future refinements to the control logic, higher throughput implementations can enable human-sized genome and epigenome mapping applications.

## Introduction

Precisely mapping the location of sequence motifs within individual double-stranded DNA (dsDNA) molecules in heterogeneous samples is central to a wide-range of genomics applications.^1^ One candidate approach for multiplexed molecular feature mapping measures ionic current modulations that arise when a dsDNA is electrically driven through a solid-state nanopore. Solid-state nanopores have sufficient sensitivity to detect a wide-range of features bound to translocating dsDNA and ssDNA, including anti-DNA antibodies, ^2^ streptavidin,^3,4^ transcription factors,^5^ histones,^6^ and peptide nucleic acids.^7,8^ They can also detect structural features involving nucleic acids such as ssDNA versus dsDNA regions,^9^ DNA-hairpins,^10,11^ multi-way DNA junctions^12^ and aptamers.^13,14^ Compared to protein pores, solid-state pores can sense a wider range of analytes due to their configurable pore diameter, which can be tuned to optimize sensing of a particular bound DNA feature.^15^ While robust technology platforms already exist for high-throughput optical mapping based on detection of fluorescent labels,^1^ solid-state nanopores offer an electrical readout with potentially higher resolution than the 1000 bp resolution limit of optics. Nanopore resolution is limited by the nanopore-spanning membrane thickness, which in principle can achieve *<* 1 nm when implemented with atomically thin materials.^16^

Despite this promise, solid-state nanopore feature mapping along dsDNA has been demonstrated only in experiments that contain pools of identical molecules or simple mixtures of known sequence,^3,10,11^ including in our own work.^17,18^ Exploiting solid-state nanopores in medical or industrial genomics applications will require a non-trivial scaling of current solid-state nanopore sensing to analyze complex samples. Such samples consist of heterogeneous mixtures of ∼10-100 kbp dsDNA fragments drawn from random locations on genomes larger than one million base pairs (Mbp) and up to human-genome scale (3,200 Mbp). Scaling of the current construct-level experiments with solid-state nanopores to larger genomes will require an enabling methodology that is capable of aligning random molecular fragments to a reference genome or to other reads for constructing contigs for genome-wide assembly.

There are three fundamental obstacles to achieving high-quality genome-scale alignment of solid-state nanopore data. Firstly, molecular folding during dsDNA translocation interrupts the linear ordering of molecular features, so a clear map cannot be established. For example, more than 60% of 48kb lambda DNA passes through a ∼10 nm diameter pore in a folded configuration. ^19^ While smaller pores that promote dsDNA linearization can be made *in situ* with additional circuitry and logic, ^20^ features bound to the dsDNA would not routinely pass through such pores. Secondly, high molecular fluctuations during translocation introduce significant random error which inhibits detection of features and alignment. Construct level barcoding experiments performed with solid-state nanopores suggest a broad distribution in translocation times;^3,10^ this is believed to arise from both fluctuations in initial molecule configuration and diffusion processes arising during translocation.^21^ Thirdly, genomic alignment requires converting barcodes from the translocation time domain data (microseconds to milliseconds) to units of genomic distance (bp). This conversion is non-trivial because it requires knowledge of the translocation speed. While the translocation speed can be obtained by assessing the translocation time between labels of known separation (e.g., using customized dsDNA calibration molecules with known label patterns^22^), this is problematic when working with complex samples containing fragments of varying length and sequence motif label patterns. In particular, there is evidence from experiments,^22,23^ Brownian dynamics (BD) simulation and tension-propagation models^24^ that the translocation velocity is non-uniform and can depend in a complex way on molecule size. Therefore, direct measurement of speed on a single molecule basis is required to enable mapping distance estimation.

We have demonstrated that dual-pore devices combined with active control logic implemented on a Field Programmable Gate Array (FPGA) can systematically address these challenges using ∼ 20 nm diameter pores. Such pores are compatible with scalable lithography fabrication methods^25,26^ but are too large to prevent folding in single-pore configurations. A dual-nanopore device features not one but two pores.^17,27–30^ To realize greater functionality, dual-nanopore devices have been developed that permit independent biasing and current sensing at each pore.^29,30^ In such dual-pore devices, if the two pores are co-located within ∼2 microns or less, it is possible to achieve a dual-capture event where different regions of a single dsDNA molecule simultaneously translocates through both pores.^27–30^ With the addition of active control logic, the two pores can exert opposing electrophoretic forces on the DNA, leading to a tug-of-war state^17^ that achieves a controlled and reduced-speed translocation. In addition, exploiting independent sensing at the pores, the time-of-flight (TOF) of a molecular feature translocating between the pores can be accessed.^17^ The TOF, using the known pore-to-pore spacing, can be used to deduce the local translocation velocity, which in turn can be used to calibrate the blockade profile in units of genomic distance. Lastly, we have supplemented tug-of-war control with active bi-directional control.^18^ Using FPGA logic to repeatedly change the direction of molecule motion during a tug-of-war event, we can induce a back-and-forth multi-scanning or “flossing” of each co-captured DNA. Multi-scanning enables acquisition of multiple reads of the same molecule so that random errors can be minimized through aggregation, while also removing folds to linearize the molecule.

Here, using our FPGA-based dual-pore platform, we take an important step towards scaling solid-state nanopore technology to analyze complex genomes, mapping a complex mixture of molecules to the 4.6M• bp genome of *Escherichia coli* (*E. coli*). To achieve this, we first decorated high molecular weight input dsDNA with molecular features formed from incorporating oligodeoxynucleotide overhangs^31^ at sites established by a nicking endonuclease. These installed features, which we refer to as “tags”, give rise to a strong localized current blockade while minimizing pore interaction for facile tag detection during DNA translocation. Next, our existing dual pore multi-scanning technique was enhanced by adding the capabilities of progressively zooming out on molecules during scanning, and of molecule recapture after exiting tug-of-war. With the recapture capability, it becomes feasible to maximize the portion of barcode scanned for each captured molecule. Time domain to genomic distance calibration was then performed using *in situ* velocity data captured using each tags local TOF data. We developed computational tools to combine extracted single fragment barcodes from *E. coli* genomic DNA into a consensus alignment, which when combined with molecule re-capture increases the average alignment performance. The presented methods comprise the most advanced single-molecule control system ever developed.

## Results and Discussion

### Experimental setup

Dual-pore DNA capture and multi-scanning experiments are performed using our previously described setup and devices.^17,18,30^ The DNA molecules were first labeled by placing 60 nucleotide long oligodeoxynucleotide (OdN) tags at the recognition sites established by nicking enzymes (Methods, Conjugation of oligodeoxynucleotide tags). The OdN tags create sharp current blockade signals that can be easily distinguished from the baseline DNA blockade level during co-capture events. The DNA is then introduced in the top reservoir referred to as the “common chamber” of the borosilicate glass chip containing the nanopores and two opposing nanofluidic channels. A dual-pore flow cell requires 7 *μ*L of 1 ng/*μ*L DNA sample, and DNA longer than ∼10 kb to ensure reliable co-capture and active control capa-bilities.^17^ The nanopores are 20-30 nm in diameter and are formed via focused gallium ion beam milling in a ∼30 nm thick silicon nitride membrane at the point of closest approach of the two channels; the nanopores thus form the fluidic gate between the nanofluidic channels and the common chamber.^30^ The pores are placed ∼500 nm apart so that a single dsDNA molecule greater than a few kbp in size can span the inter-pore distance and simultaneously thread through both pores. Critically, our design permits independent control of voltage biasing as well as independent acquisition of ionic current signal at each pore. This allows simultaneous detection of DNA in each pore while separately adjusting the voltages applied across each pore. Active logic implemented on an FPGA adjusts the voltages in response to changes in the ionic current signal at either or both pores, leading to flexibility in control protocol design.^17,18^ The instrumentation and alignment algorithms explored here were first benchmarked using *λ*-DNA with OdN flaps placed at a superposition of sites for the enzymes Nt.BbvC1 and Nb.BssSI. The challenge of genome scaling was then demonstrated using DNA extracted from *E. coli* with OdN flaps placed at Nb.BsrDI sites (Methods, Isolation of genomic *E*. *coli* DNA), as a benchmarking exercise that was compatible with the throughput of the current non-arrayed dual-pore device implementation.

### Flossing with zoom-out mode and repeated DNA captures

The FPGA is configured to enhance the probability that a molecule transiting the dual-pores will be captured in a tug-of-war state.^17^ Once captured in tug of war, the biasing at the pores are set to move the DNA in a controllable direction, i.e., with DNA moving in the direction of the pore with the larger channel-side positive voltage bias. The FPGA is also used to dynamically change the voltage bias magnitude so that the molecules direction is changed, with multiple sequential changes producing multiple back-and-forth interrogations of the same molecule in what is termed “DNA flossing.”^18^ During DNA flossing, the voltage bias at one nanopore is changed in step-wise fashion each time a change in direction is triggered, while holding the bias at the second pore constant. We will refer to the pore where the bias changes as pore 1 and the pore with constant voltage as pore 2. DNA motion from pore 1 to pore 2 is referred to as “left to right” motion (L-R); DNA motion in the opposite sense from pore 2 to pore 1 is referred to as “right to left” motion (R-L). In this study, flossing voltages at pore 1 are 150 mV for L-R motion and 650 mV for R-L motion, and remains 300 mV at pore 2.

In our flossing implementation, a co-captured molecule is initialized with motion in the L-R direction, and the controller is programmed to switch scan direction to R-L after a preset number of tags are detected in the pore 2 current. Similarly, the controller switches scan direction back to L-R once the same number of tags had been observed in pore 2, leading to a full cycle of the flossing motion. Triggering scan-direction changes based on a finite number of tag detections introduces a set of challenges when working with DNA fragments of unknown length and that contain an unknown number of tags. For such DNA fragments, for example, if the preset number of tags is too small, we will achieve a large number of scans but fail to scan other portions of the molecule, while also spending too much measurement time on a small region of one molecule. On the other hand, if we set the preset too large then we risk having the molecule disengage from dual-pore capture during the first scan event, preventing the cyclic flossing function altogether. In order to achieve a balance between maximizing the scan number and scanning a sufficiently large portion of the barcode, we have developed an adaptive strategy in which the controller will “zoom out”, iteratively increasing the preset number of tags after a finite number of successful flossing cycles.

For a newly co-captured molecule from the top chamber, the zoom-out controller starts by scanning for two tags to trigger changes in scan direction, as previously described and demonstrated .^18^ After a preset number of scans (nominally 4 scans L-R and 4 scans R-L), the tag count is increased by one and the multi-scanning continues. A depiction of this zoom out process with representative data is shown in Figure 1 for the 4-tag-count and 5-tag-count stages, showing 2 representative scans for each scan direction. The zoom out process continues until the tag count reaches a maximum that is set by the user, nominally set to 8 or more tags depending on the anticipated tag density range. The molecule eventually exits co-capture and flossing for a variety of reasons, including those previously described^18^ such as drift in the motion of the molecule during flossing and tags that go undetected and thus uncounted (Methods, Tag Calling). Molecules can also exit co-capture when the total tag density is lower than the maximum tag count.

**Figure 1:**
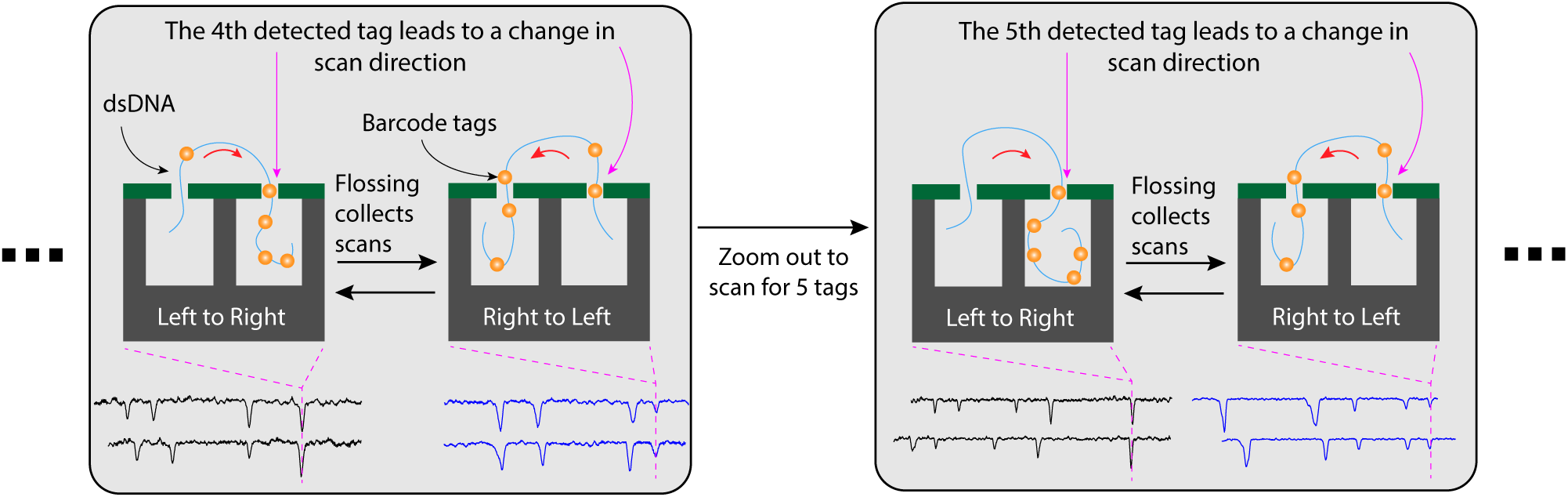
A cartoon depiction and recorded data showing an example of the “zoom out” control function that occurs during DNA flossing. The dsDNA with barcode tags are depicted as a blue line with orange circles, respectively. In the example shown, the tag-counting limit is 4 (left shaded box) and a captured molecule is scanned left to right (L-R) until the 4^th^ tag is detected in pore 2, and then right to left (R-L) until the 4^th^ tag is detected in pore 2. Once a preset number of scans are collected, the controller zooms out by performing the same logic but waiting for the 5^th^ tag before changing direction (right shaded box). The process is repeated with the tag-counting limit increased up to a user defined limit (e.g., 8 tags) and/or until the molecule exits co-capture. The pore 2 signal traces are representative scans from a co-captured fragment from *E. coli*.

In addition to the zoom out function, we further enhanced the logic to automatically recapture the molecule after exiting from the tug-of-war state (Figure 2(a)). Exit in either scan direction leads to the molecule being located in one of the nanofluidic channels, and the recapture logic is different depending on the direction of DNA motion when exit occurs. In the event of a R-L exit, the molecule is in the channel below pore 1 (Figure 2(a), State 5a), and the logic restarts the process of achieving a tug-of-war state using precisely the same logic sequence when a molecule is captured initially from the top chamber.^18^ In the event of a L-R exit, the molecule is in the channel below pore 2 (Figure 2(a), State 5b) and the logic implemented is modestly more complex. First, pore 1 voltage is set to 0 mV and the voltage is reversed at pore 2 to re-capture the DNA from the channel back through pore 2. Note that recapture into a pore from the channel has a high probability due to the influence of voltage along the length of the nanofluidic channel.^17^ Next, after the DNA is fully ejected through pore 2 and into the common chamber, pore 2 voltage is set to 0 mV and the pore 1 voltage is turned back on to recapture the DNA through pore 1 and to return to State 1 in Figure 2(a). The times scales of time-of-flight recapture (pore 2 to pore 1) are fast when recapture occurs,^30^ and are thus time bounded in the logic by a waiting period of 1 s to maximize the probability of capturing the same molecule in pore 1 that exited pore 2, rather than a new molecule from the common chamber. By contrast, the time to capture a new molecule in pore 1 from the common chamber occurs on a time scale that is comparatively slower than the pore 2-to-pore 1 recapture process, i.e., at the 1 ng/*μ*L concentrations used. For the merged *E. coli* experiment data, for example, the exponentially distributed time-to- capture of new molecules from the common chamber had a mean 10.6 s and 10^th^-percentile of 1.1 s, all of which are slower than the maximum wait time of 1 s for recapture.

**Figure 2:**
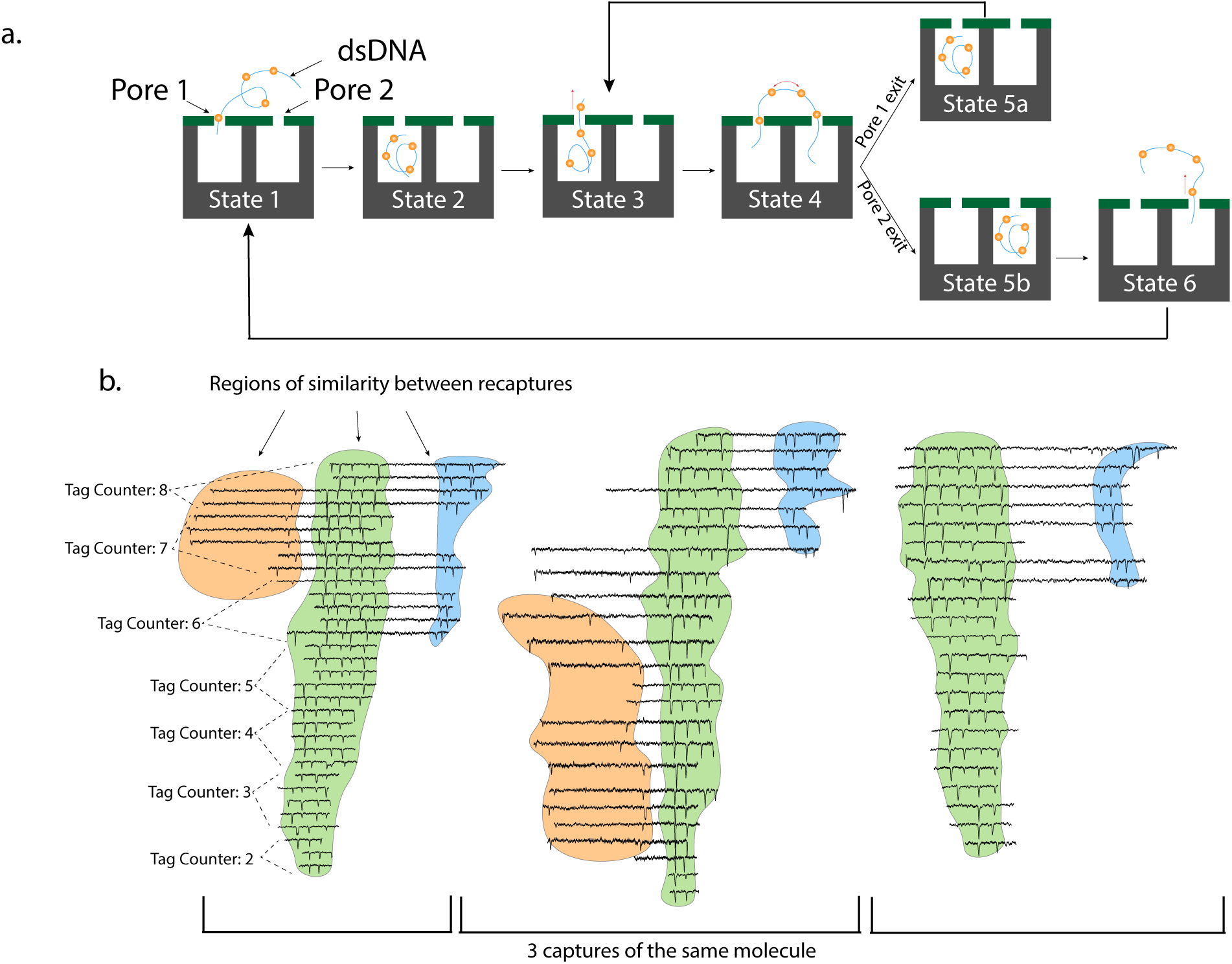
A cartoon depiction and recorded data showing the collective set of single-molecule data that DNA flossing with recaptures can produce. **a**. The cartoon shows the recapture procedure, with state 4 comprising the flossing with zoom routine. When the DNA is lost to channel 1 (pore 1 exit, state 5a) the controller triggers tug-of-war and flossing without searching for another molecule from the bulk common chamber. When the molecule is lost to channel 2 (pore 2 exit, state 5b), the controller moves the DNA from channel 2 into the area directly above the two nanopores, priming it for re-capture (state 1) and restoration of flossing. **b**. Representative traces from flossing with zoom on a single molecule (each vertical panel) comprising a total of three captures. The tag-counting limit used during zoom-out is reported for the left-most flossing event. The molecule was a fragment from *E. coli* and signals are as measured at pore 2 in L-R direction. The scans were produced chronologically from bottom to top in each panel. Drift in the molecule during zoom out can occur due to missing tags in the counting logic, which creates a frame shift in the region of the molecule being scanned.

Figure 2(b) gives an example of a subset of scans from three recaptures of the same molecule. Each flossing event was recorded from bottom to top, with the first flossing event annotated (left side) with the increasing tag-counting limit that was used during zoom-out control. Recapture events commonly show similarity in the barcode patten, as observed for the shaded regions of the signal traces in Figure 2(b). Also observable are the stochastic variations in the signal in the form of variable tag amplitudes and variable inter-tag durations. Variations in tag amplitude can lead to imperfect detection and counting of tags during flossing, which in turn contributes to drift in the DNA motion during flossing and thus drift in the scanned region of the molecule over sequential scans. The variation in inter- tag durations is visibly larger where the tags are spaced farther apart, e.g., for the time gap between the right most tag in the green shading and the left most tag in the blue shading in Figure 2(b). Variable inter-tag durations can contribute to larger spread in predicted distances. However, distance estimation variation between *a priori* unknown tag locations is ameliorated here by using direct measurement of molecule velocity. Specifically, a differentiating advantage of the dual-pore system is that the velocity of individual tags can be directly measured on a per-scan basis,^18^ which enables inter-tag distance estimation for each scan as described in the next section. Our direct velocity measurement requires no calibration, and can be done directly for any tag detected at both pores. This advantage means that while regions can have variable scan speeds that create variable inter-tag times, direct knowledge of velocity can be exploited to produce more consistent distance estimates, as suggested in related studies.^32^

### Alignment of single scans to a reference map

A scan is a single pass of a region of a dsDNA molecule moving through the dual-pore sensors in either the L-R or the R-L direction. For a scan to be align-able to a reference map, it must first have electronically detectable tags. A reference map is a list of genomic positions generated by digesting the reference genome *in silico* by one or more nicking endonucleases (Figure 3(a)). Our objective is to align the sequence of tags observed in the scan to the positions in the reference map by matching the distances between observed tags to the known number of base pairs between nicking sites. This requires estimating the distances between observed tags in genomic coordinates. To estimate distance in genomic coordinates requires two conversions: 1) converting the time period observed between any two tags into a distance estimate in spatial linear coordinates (units of nanometers); and 2) converting the spatial distance estimate into a distance estimate in genomic coordinates (units of base pairs). The first conversion from inter-tag time to inter-tag linear distance utilizes the known linear distance between the dual nanopores (Methods, Estimating scan velocity and linear distance between adjacent tags). The second conversion requires consideration of the stretching behavior that occurs between the pores as a result of the tug-of-war forces, which is expected to be asymmetric in the two scan direction since higher tug-of-war voltages are used R-L than L-R. The details of these two sequential conversions are described next.

**Figure 3:**
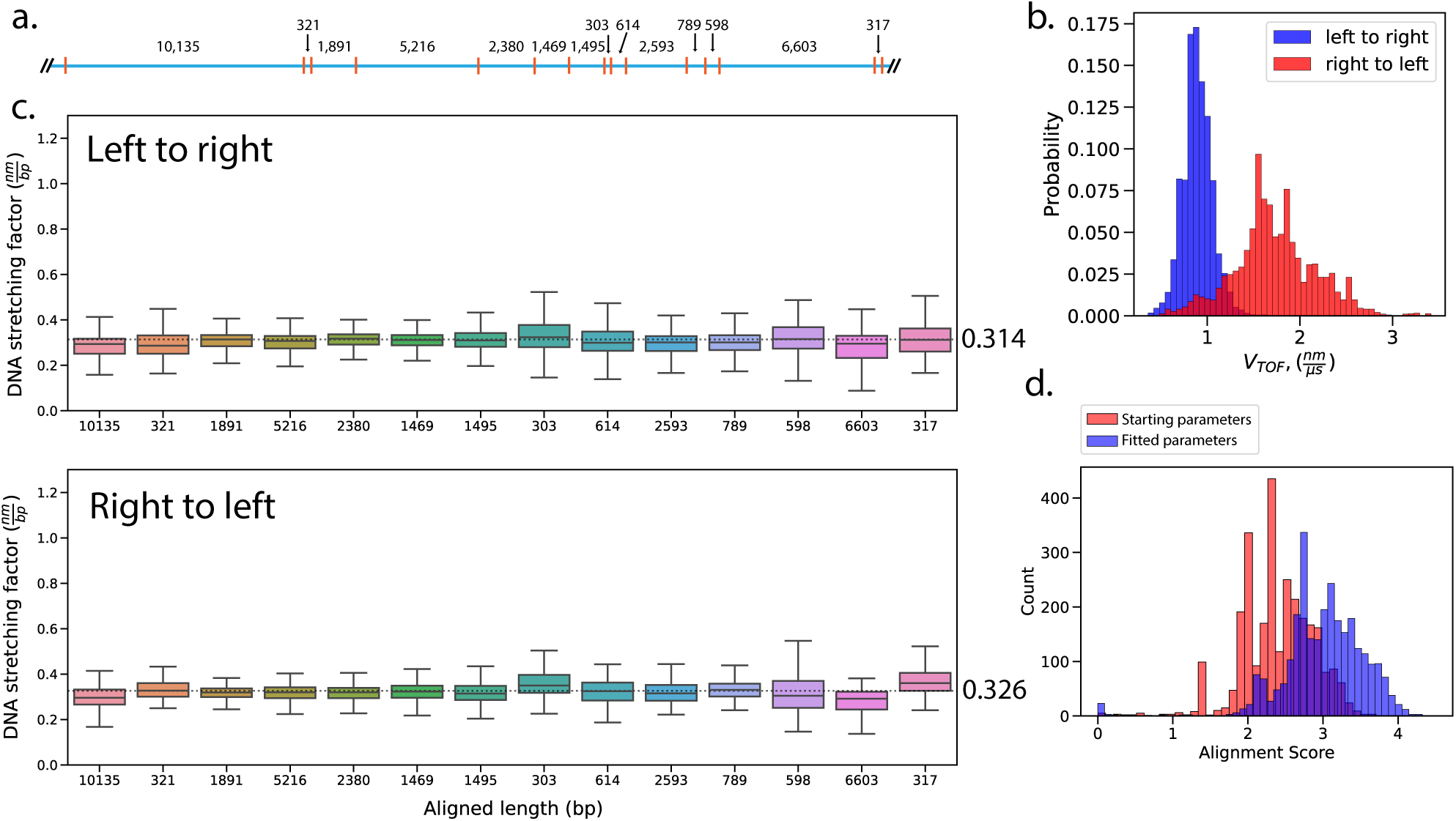
Performance of alignment of individual scans from flossing data using a model 15-tag *λ*-DNA. **a**. Visual description of the 15-tag phage *λ*-DNA test molecule. Orange hashes indicate nicking sites from the Nt.BbvC1 and Nb.BssSI enzymes. **b**. Histogram of the measured *V*_TOF_ for L-R (blue) and R-L (red) scanning directions, with data drawn from 7,711 tags within 1,724 scans L-R, and 3,716 tags within 1,074 scans R-L. **c**. DNA stretching factor (nm/bp) for the different intervals on the *λ*-DNA test molecule for L-R (top) and R-L (bottom) for the same scans used in (b). We observe small variations around the weighted average, 0.314 ± 0.05 nm/bp and 0.326 ± 0.05 nm/bp for L-R and R-L scans, respectively (black dotted line, annotated to the right). **d**. Histogram of single-scan alignment scores for the initialized and fitted model parameter. The scores improved by using the direction-specific models and the optimized probability distributions.

To estimate the spatial linear distance *D*_*n,m*_ between the *n*^th^ and *m*^th^ tags detected in a scan, we first identify the positions of the *n*^th^ and *m*^th^ tags in the time-domain, which is accomplished by identifying the times associated with the peak current attenuations for the pair. To convert from the time-domain to units of linear distance, we measure the velocity of tags during flossing and create a velocity profile that linearly interpolates between these values. Briefly, when a tag is detected at each pore, the time required for the tag to traverse the inter-pore distance is obtained, and is referred to as the tags time-of-flight (TOF). The inter-pore distance divided by the tag TOF yields the tags time-of-flight velocity *V*_TOF_, and a velocity profile of the DNA chain itself during the scan is generated by linearly interpolating between sequential *V*_TOF_ values (Methods, Estimating scan velocity and linear distance between adjacent tags). The linear distance *D*_*n,m*_ between any two sequentially detected tags *n* and *m* is then computed by simply integrating the scan velocity profile over the inter-tag time period.

The average of each scans velocity profile can be examined to assess the DNA chain speed distribution across scans, and in each scan direction. In our experiments on *λ*-DNA, we estimated the mean L-R velocity to be 0.89 nm/*μ*s, which is ∼2X slower than the mean R- L velocity of 1.73 nm/*μ*s (Figure 3(b)). Larger voltages create larger DNA velocities through a nanopore, and in the dual-pore setup the net voltage difference defines the net force on the molecule. Therefore, the 2X higher observed velocity going R-L is likely a consequence of the 2X larger net force: 300 mV R-L (600 − 300 mV) vs. 150 mV L-R (300 − 150 mV).

Once linear distance estimates between tags are computed for a given scan, the second conversion requires converting the spatial distance estimates (nm) into a genomic distance estimates (bp). The core of our alignment method is a probability distribution modeling the stretching of DNA under forces exerted by the dual-pore system. Let *μ* be the DNA stretching factor (nm/bp), and let *G*_*i,j*_ be a genomic interval distance (bp) found in the reference and corresponding to nicking site indices *i* and *j*. With a given constant stretching factor *μ* presumed for a given scan, the alignment problem is to find the optimal placement of the scan data {*D*_*n,m*_} to the reference data {*G*_*i,j*_} such that 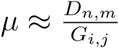, and while searching across the entire reference and accommodating for false-negative and false-positive tags. To achieve this, we first model the probability of observing a given DNA stretching factor by a normal distribution 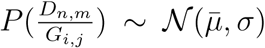 with mean 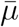, and standard deviation *σ*. The probability density is

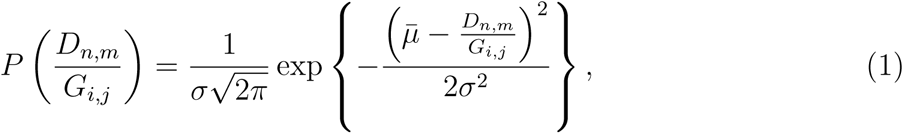

and we convert this into a score model by taking the log of the probability density. The score model allows us to quantify how well a measured set of distances between tags matches a corresponding set of genomic intervals between nicking sites on the reference. Previous studies have shown B-form DNA to have a stacking height of approximately 0.34 nm/bp.^33,34^ In order to make an initial estimate of 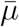 and *σ* we initialized our model with 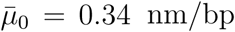 and *σ*_0_ = 0.12 (Methods, Scan-space alignment). This score model is then built into an alignment algorithm similar to Smith-Waterman local alignment^35^ that accounts for false-negative and false-positive tags by the tag calling algorithm (Methods Description of alignment algorithm).

The alignment method was applied to experimental data and a reference map based on *λ*-DNA with incorporated tags at 15 nicking sites (Methods Conjugation of oligodeoxynucleotide tags). This relatively simple reference map permitted alignments to be manually inspected for correctness. Scans collected from 27 dual-pore chips were filtered (Methods, Data processing and filtering), resulting in 2,897 individual scans within 889 co-capture events composed of 409 individual molecules. The distribution of fitted stretch factors were obtained separately for L-R and R-L directions, and stayed consistently near their averages of 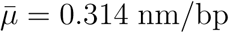 for L-R direction and 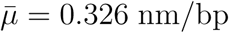 for R-L direction across the length and varying inter-tag distances of the *λ*-DNA molecule (Figure 3(c)). These values indicate that the DNA, while strongly extended, is not 100% stretched as a semi-flexible chain at the tug-of-war voltage values used.^17^ The modestly higher stretching coefficient for the R-L direction is presumed due to the higher tensile stretching forces than L-R.

The stretching factors showed high variability on a per site basis (Figure 3(c)), as indicated by high standard deviations: *σ* = 0.25 nm/bp for L-R and *σ* = 0.34 nm/bp for R-L. To obtain an improved estimate, we performed a weighted average where the weights are the probability of the aligned segment using the initial values of 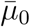 and *σ*_0_. The probability-weighted average suppressed the influence of low probability outlying pair measurements and improved the per-site estimate variability. The weighted estimates yielded 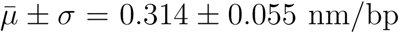 for L-R direction and 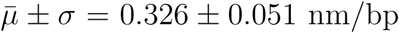 for R-L direction (Table 1). Upon iterative realignment of the *λ*-DNA results using these refined values of 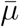 and *σ* for L-R and R-L directions, we observed an increase in alignment score of 6.1% (Figure 3(d)).

**Table 1:** Parameters in model Equation (1) after weighted averaging.

| | number of scans | $\bar{\mu}$ | $\sigma$ |
| --- | --- | --- | --- |
| left to right | 1724 | 0.314 | 0.055 |
| right to left | 1074 | 0.326 | 0.051 |

### Tag resolution is at least 150 base pairs

One advantage of electronic nanopore based measurement over optical detection is the potential to resolve smaller distances.^3^ To test the resolution of our dual-pore instrument, we engineered specific *λ*-DNA reagents possessing closely spaced tags. Cas9 nickase was used to install 60 nucleotide ssDNA tags separated by 150 bp (Methods, Construction of 150 base pair tag-pair DNA). We observed 23 molecules with two distinct spikes due to the two tags (Figure 4(a)). We calculated the tag-pair resolution *T*_res_ using a formula from liquid chromatography

**Figure 4:**
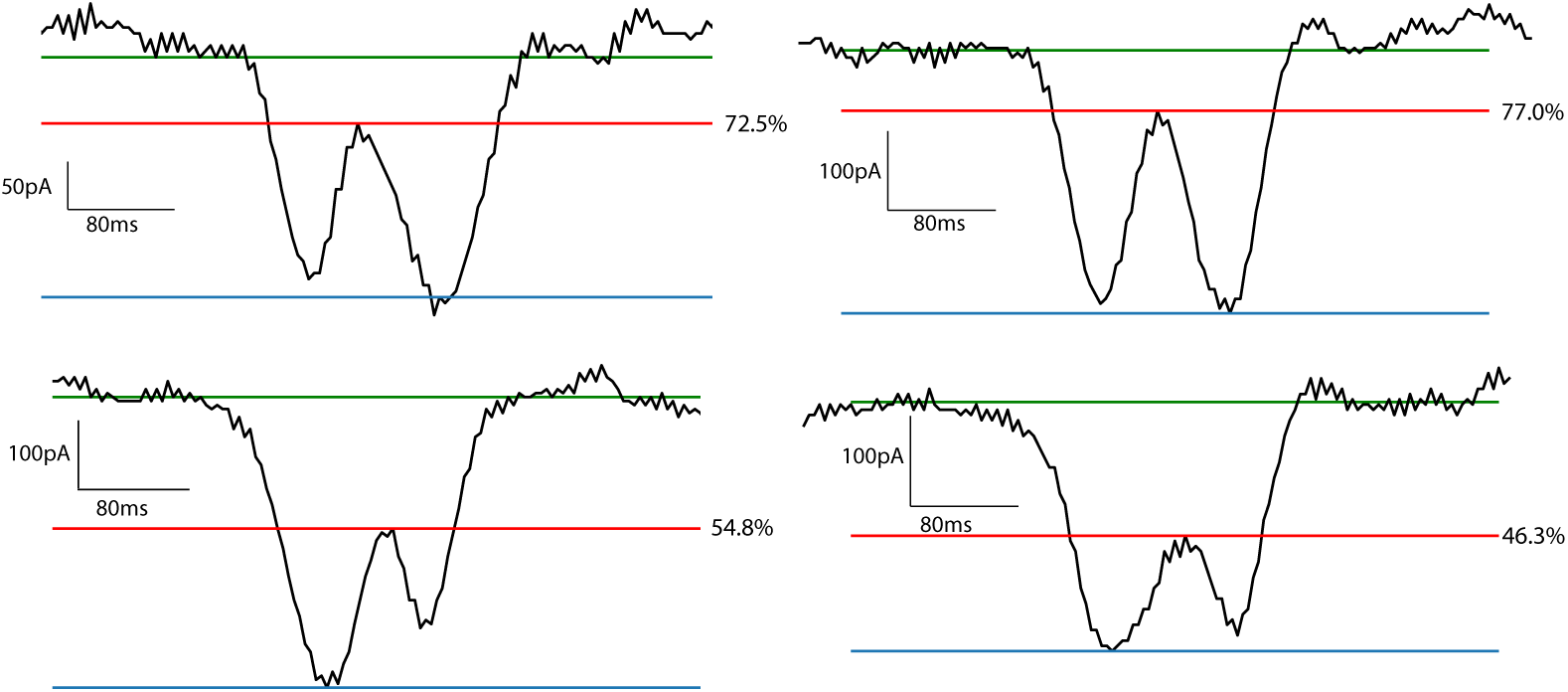
Representative current traces of 60 nucleotide tags separated by 150 bp. Green, blue, and red lines show the estimated tag-free DNA baseline current, the minima current attenuation created by one of the two tags, and the restoration percentage line (with value reported), respectively.

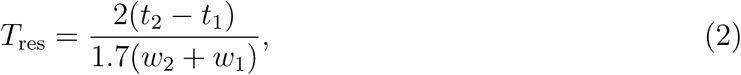

where the first and second tags have peak minima at times *t*_1_ and *t*_2_ and peak time widths at half-minimum depth of *w*_1_ and *w*_2_, respectively. We also measured the percentage of the restoration of the signal between the two peaks. This “restoration percentage” is the percentage that the signal achieves from the minima current attenuation up to the tag-free baseline signal. Statistics from the tag-pair resolution values from Equation (2) and the corresponding restoration percentage values are reported in Table 2. While the signals for tag pairs are resolvable at 150 bp here, optical mapping has a 50% chance of resolving two fluorescent reporters within 1000 bp.^36^ Additionally, future instrumentation at 2X higher bandwidth than the 10 kHz low-pass Bessel filter used here should provide 2X sharper peak resolution with an acceptable increase in high frequency noise, which will further increase the resolution score and permit exploring the detection of tag pairs closer than 150 bp.

**Table 2:** Tag-pair resolution and restoration percent for 23 molecules.

|  | Restoration Percentage | Tag-Pair Resolution |
| --- | --- | --- |
| mean | 58.59 | 0.77 |
| std | 10.74 | 0.23 |
| min | 34.34 | 0.48 |
| max | 77.06 | 1.17 |

### Generating consensus alignments from multi-scan flossing events

A flossing event produces a significant quantity of information distributed over multiple scans. This information needs to be assembled into a best estimate of the overall pattern of tags present on the analyzed fragment. To this end, we developed an algorithm that takes a set of single scan alignments from a flossing event and assembles these alignments into an overall consensus alignment.

Our algorithm first aligns the individual scans to the reference map using our alignment algorithm with a tuned Gaussian scoring function. To assemble the alignments into a consensus we consider the problem in a graph theoretic sense. Let *𝒢* be a graph with vertices *v*_*i*_ ∈*𝒱* and edges *e*_*ij*_ ∈*ε* connecting vertices *v*_*i*_ and *v*_*j*_. Each vertex, *v*_*i*_, represents a nicking site at position *i* in the reference map (in bp). Each alignment corresponds to a sequence of tuples *A* = [(*a*_0_, *s*_0_, *d*_0_), (*a*_1_, *s*_1_, *d*_1_), …, (*a*_*k*_, *s*_*k*_, *d*_*k*_)]. The quantity *a*_*k*_ represents an aligned pair *a*_*k*_ = (*n, m*) ◊ (*i, j*), the notation indicating that the *n*^th^ tag in the scan is aligned to the nicking site at position *i* in the reference and likewise that the *m*^th^ tag in the scan is aligned to position *j* in the reference. In addition, each aligned pair *a*_*k*_ has an associated score *s*_*k*_ and measured distance *d*_*k*_. The score is calculated in Equation (1) and using the parameters 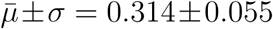 for L-R data and 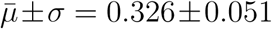 for R-L data (Table 1). We start by initializing *ε* = ∅ (i.e. no alignments in the set) and proceed with graph construction by iteratively adding each alignment to the graph. For each aligned pair *a*_*k*_ we add an edge connecting vertex *v*_*i*_ and *v*_*j*_ with the edge weight equal to *s*_*k*_. If an edge already is present from a previous added alignment, we simply add *s*_*k*_ to the weight corresponding to that edge. We also maintain a mapping, *e*_*ij*_ → *D*_*ij*_, of the measured distances of the aligned pairs for each edge in the graph. Following graph construction, the graph is pruned to remove edges with scores below zero. The consensus alignment, 𝒜, is reported as the maximum scoring path from a start vertex to an end vertex. Lastly, the measured distance for a given interval in the consensus alignment is the average of the measured distances for the edge connecting the interval 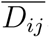.

To convey the process of generating consensus alignments from multi-scan flossing events conceptually, a synthetic reference and examples of synthetic scans are shown in Figure 5(a). The concept shows the need for at least two scans to detect conflicts, and at least three scans to resolve conflicts (site 3 vs. site 4). The concept also shows that the resulting consensus can have missed tags (site 3) and missed regions (sites 8-10). The results of applying the process to one representative multi-scan flossing data set from *λ*-DNA in which all 15 tags were detected is shown in Figure 5(b). After removing 1% of outliers, the performance of the consensus estimates from all *λ*-DNA flossing data is shown in an error histogram in Figure 5(c).

**Figure 5:**
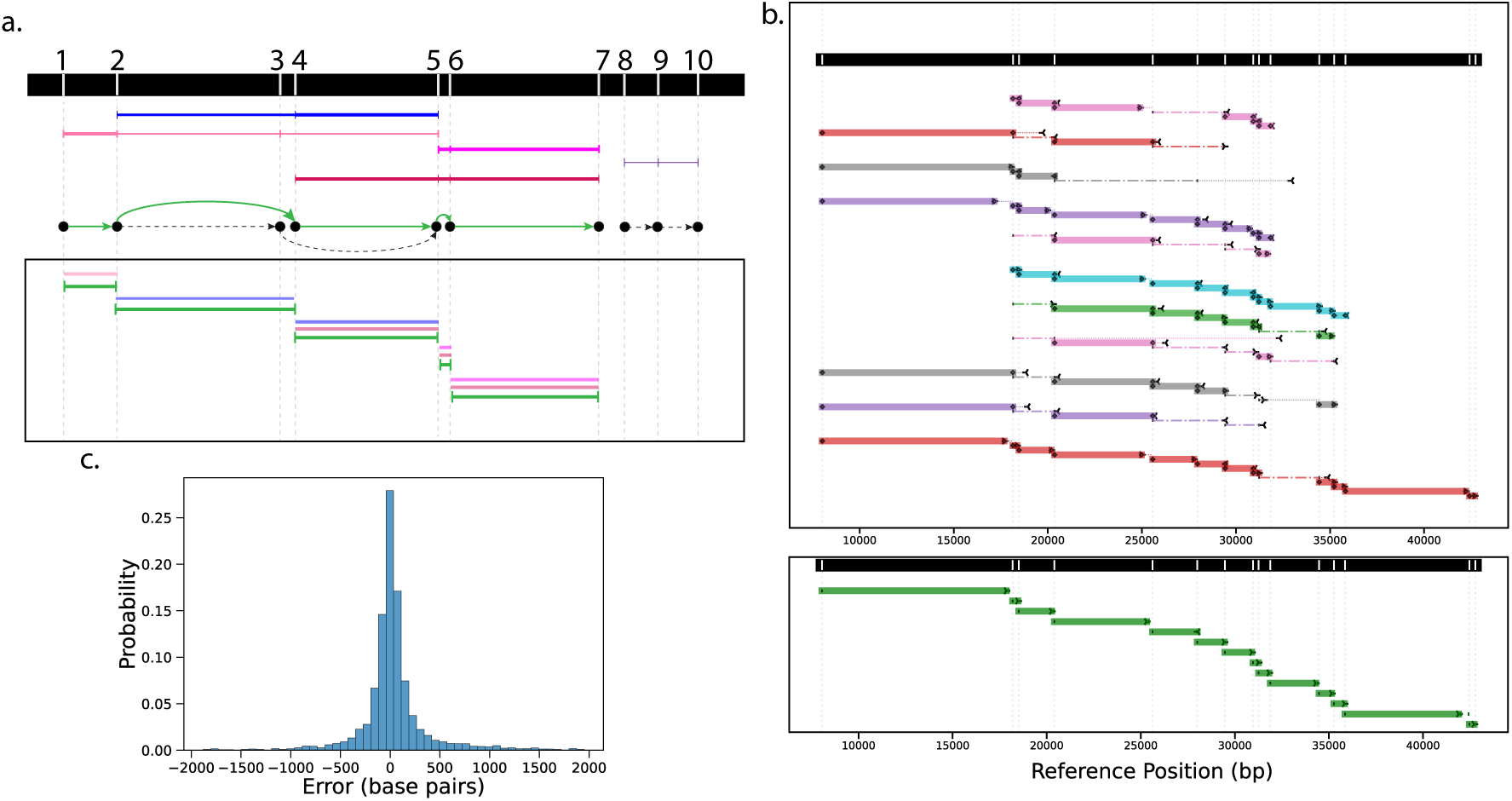
Concept and recorded data show how consensus alignments are produced from flossing event data. **a**. A fictional reference map with ten tagged sites, and five conceptual aligned scans with line thickness proportional to the relative fitting score. The four scans spanning sites 1-7 (colors: blue, pink, magenta, and red) are used to identify the highest scoring directed graph path (1, 2, 4, 5, 6, 7) (solid green lines with arrows), while the scan spanning sites 8-10 (purple) is low scoring and not connected to the rest of the graph and is thus ignored. An alternative path (1, 2, 3, 5, 6, 7) (inserted dashed lines) would require ignoring the more probable and scan-supported 4-5 path transition and thus produces a lower score. The consensus alignment (box, green lines) shows the supporting scan intervals by color that are averaged to generate the consensus distance estimates. **b**. Interval plots of individual scan alignments (upper box) and resulting consensus alignment (lower box) from a single recorded *λ*-DNA flossing event. Each aligned pair of tags are shown as colored bars if sufficiently high scoring and thus used in the consensus, or as dashdotted lines if low scoring and not used in the consensus. Bars and lines of the same color are from the same scan. The bar or line length indicates the estimated number of base pairs for that interval, with a vertical offset of adjacent intervals used to help visualization. Dotted lines are inserted where length estimates appreciably deviate from their assigned reference values, with arrows to demark the endpoint of estimates and the direction of the assigned reference endpoint. The consensus (bottom) produced estimates for all 15 sites from the first to the last tag. **c**. Consensus inter-tag alignment length error histogram (Trimming 1% outliers with absolute error *>* 2 kb), comprising 3268 consensus tag-pair distance predictions across 607 captured *λ*-DNA molecules, resulting in 30 bp mean and 349 bp standard deviation for the distribution plotted and with 192 bp mean and 84 bp median absolute error.

For the *λ*-DNA results, the consensus alignment error distribution appears normally distributed with 30 bp mean and 349 bp standard deviation. Considering the absolute error as a better measure of accuracy results in a mean of 192 bp and median of 84 bp. An approximation for the 95^th^-percentile is the mean absolute error plus 2 standard deviations of the error, which is 890 bp, suggesting most of the consensus errors are below the resolution limit of optical mapping (1 kb).^36^ The value-add of consensus estimates is apparent by considering the relative performance to that of the single longest scan from each consensus molecule, as a proxy for single read data. Trimming 1% of outliers again, 3,156 high scoring estimates from the set of longest scans produced 153 bp mean and 1060 bp standard deviation for the error distribution, and 425 bp mean and 112bp median for the absolute error. Again using the the mean absolute error plus 2 standard deviations of the error results in a 95^th^-percentile error estimate of 2500 bp, nearly three times higher than the multi-scan consensus accuracy. Thus, consensus multi-scan map estimates improve accuracy and reduce the variance of distance estimates compared to single-scan nanopore data.

The process of generating consensus alignments using a multi-scan flossing molecule from *E. coli* is shown in Figure 6. These same data also annotated in Figure 2. The molecule had a total of four captures with scans organized into tracks (i-iv) in Figure 6a. For each capture, the initial scans with fewer tags align to multiple sites in the reference genome, as shown in the top lines for each track. After zooming out, subsequent scans have more tags and thus align less ambiguously to the reference, converging in their support to a locus near the midpoint of the reference genome (Figure 6b-c).

**Figure 6:**
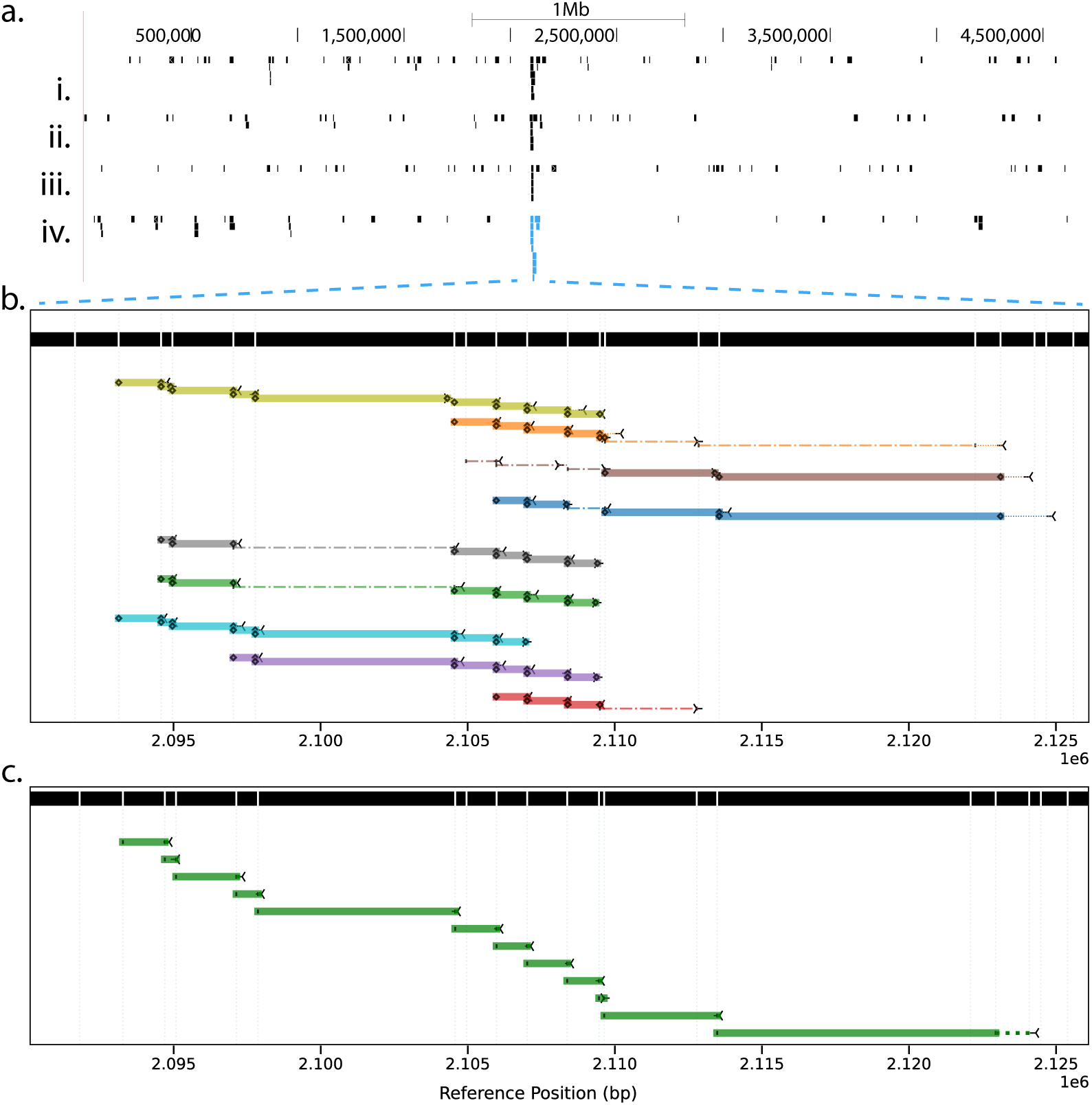
A representative consensus single-molecule alignment produced from an *E. coli* experiment. **a**. Data are from four DNA flossing captures from the same molecule. Each bar is a single scan, and each track (i-iv) is a recapture of the same molecule. Scan alignments for the fours events (i-iv) with fewer tags are spread across the *E. coli* reference genome with multiple prospective alignments, while scans with the highest tag counts achieve alignment with strong support at a unique common locus near the mid-point of the genome. **b**. Interval plots of individual scan alignments for the highlighted multi-scan data within recapture (iv). Annotations of scan details are consistent with the description in Figure 5b caption. **c**. The consensus produced comprised a total of 13 sites across the 2,075,645 to 2,178,824 locus.

### Physical genome mapping with *E. coli*

We collected 5.6 hours of dual-pore recording data from genomic *E. coli* DNA across 3 devices (Table 3). This data includes 979 single-molecule flossing events composed of 564 individual molecules. Based on our results indicating a resolution of at least 150 bp (Tag resolution is at least 150 base pairs), we modified the Nb.BsrDI reference map by merging sites that were within 150 bp of each other. We generated consensus alignments as described in Generating consensus alignments from multi-scan flossing events, requiring that the consensus alignment contain at least 3 aligned intervals, resulting in 767 consensus alignments. The data contained 247 consensus alignments with more than one scan support. The consensus alignments resolved 2,355 (68%) of the sites in the reference map including 133 inter-site intervals of less than 200 bp, with an average coverage of 3.1× over all sites (Figure 7(a)).

**Table 3:**
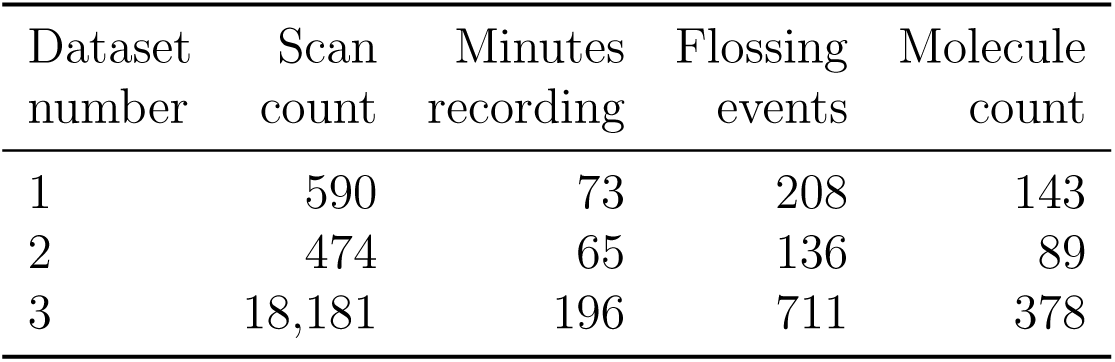
Throughput for three dual-pore devices used to generate *E. coli* data.

**Figure 7:**
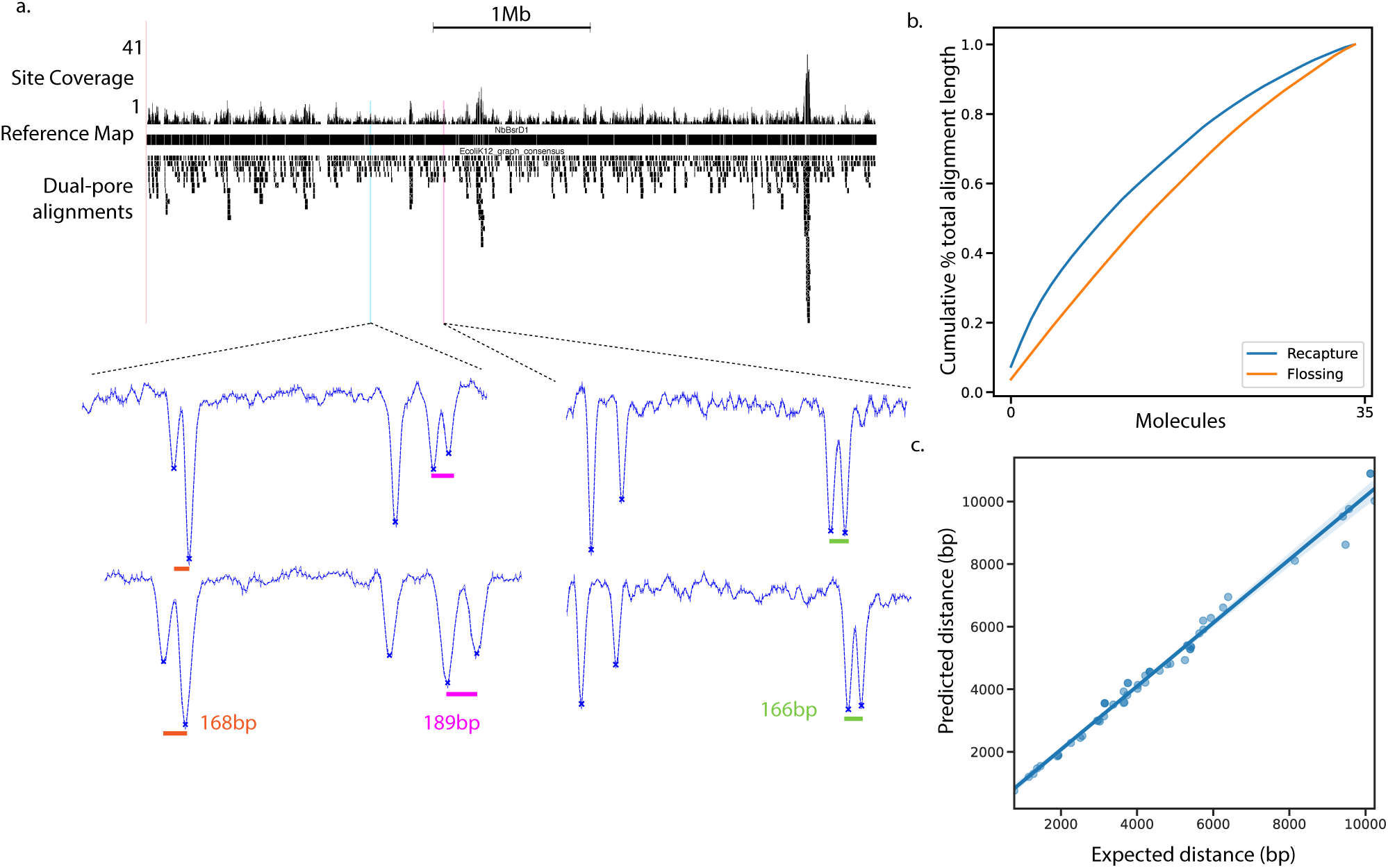
Dual-nanopore physical genome mapping of *E. coli*. **a**. (top) Microbes genome browser^38^ of the *E. coli* K12 genome and Nb.BsrDI genome map. The per site coverage, reference map sites, and consensus dual-pore alignments are shown on the top, middle, and bottom tracks, respectively. (bottom) Representative sequential scans with tags that align to distances below 200 bp in the reference genome, and bars showing the aligned distance values.**b**. Cumulative alignment length as a percentage of the total length of all alignments for recaptured molecules vs. single capture molecules (formula in Calculating N50 of alignments). **c**. Correlation of predicted and expected base pair distances for inter-tag regions spanning tandem repeats (*R*^2^ = 0.97).

To quantify alignable molecule lengths, we use the N50 (Methods Calculating N50 of alignments), noting that the length values are defined as the first-to-last tag distances and do not include the length of the molecules outside of these tags. The N50 of the consensus alignments are longer (17.4 kb) than the N50 of the longest single scan (15.8 kb) from the corresponding flossing event, since consensus alignments use the best scoring regions across all scans including the longest. We can also boost alignment lengths further by joining together, where possible, the consensus alignments resulting from multiple recaptures of the same molecule. Our dataset contained 35 molecules that were recaptured at least once after escape from initial capture. Joining recaptured molecules into a single consensus alignment increased the N50 to 21.9 kb, a 38% increase compared to the longest single-scan alignments (Figure 7(b); Methods Joining recapture alignments). Length scales of the consensus molecules including the N50 values are reported for *E. coli* and *λ*-DNA in Table 4. We also quantified the tagging efficiency as the number of observed aligned tags divided by the number of expected tag sites spanning the net consensus alignment length, resulting in 88% efficiency for the 15-site *λ*-DNA and 69% efficiency for *E. coli*.

**Table 4:**
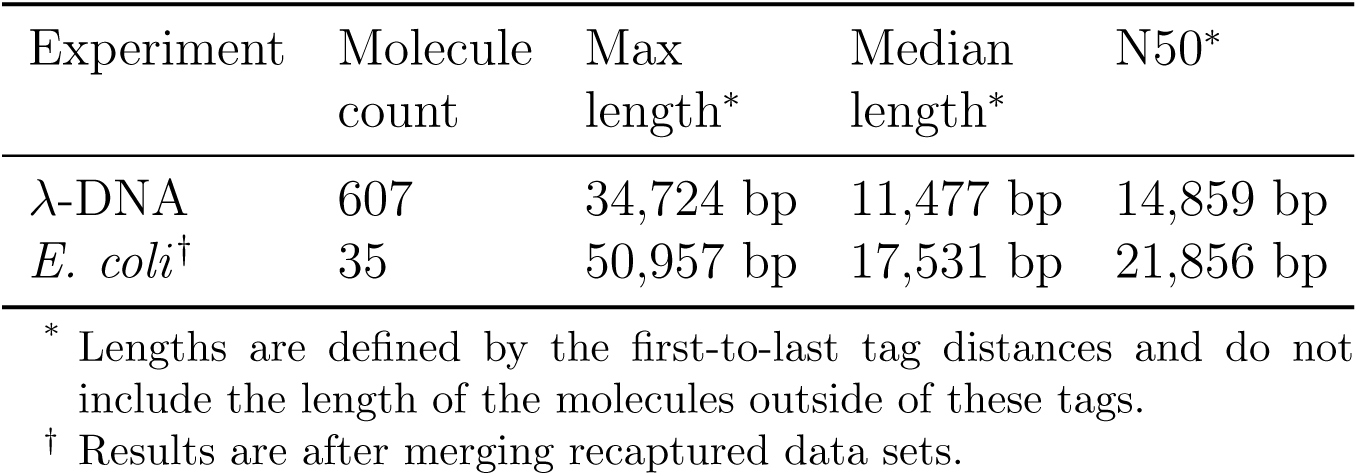
Single-molecule consensus lengths using dual-nanopore DNA flossing.

| Experiment | Molecule count | Max length* | Median length* | N50* |
| --- | --- | --- | --- | --- |
| λ-DNA | 607 | 34,724 bp | 11,477 bp | 14,859 bp |
| *E. coli*<sup>†</sup> | 35 | 50,957 bp | 17,531 bp | 21,856 bp |

\* Lengths are defined by the first-to-last tag distances and do not include the length of the molecules outside of these tags.
<sup>†</sup> Results are after merging recaptured data sets.

One potential application of physical genome mapping is to detect structural variations, including those in repetitive regions which are often difficult to resolve with short-read sequencing. We identified 93 regions of tandem repeated sequence using Tandem Repeats Finder^37^ (Supplemental File 1), to use as test regions for the accuracy of our method. We used alignments spanning the repeat region and convert their measured distance (in nm) to bp by dividing by 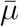 for the score model (see Alignment of single scans to a reference map). We then averaged their estimated genomic distances. Filtering to sites with at least 2 aligned molecules, our base pair length predictions showed good correlation with the expected values, *R*^2^ = 0.97 (Figure 7(c)).

## Conclusions

Our results show the first proof-of-concept demonstration of solid-state nanopore genome mapping from a heterogenous mixture of DNA molecule sizes and label densities across a mega-base scale genome. We enhanced our previous dual-pore DNA flossing approach by adding the capabilities of zooming out and repeating captures, which when combined with the presented analytical framework increases both the quality and size of consensus single-molecule alignments. We also presented proof-of-concept results estimating genomic distance in repetitive regions showing good correlation with low coverage (minimum of 2 molecules), and setting the stage for structural variation analysis with the proposed method. Our method uses relatively small amounts of input genomic DNA (7 ng / flow cell) compared to other single molecule approaches. Moreover, the nanopore technology’s purely electrical basis confers a small footprint and cost relative to optical mapping approaches. Thus, our approach has potential to produce high resolution physical genome maps with low sample input and low cost for potential field and clinical research use.

Key remaining technical challenges include data throughput and device fabrication. In particular, only 41% of the *E. coli* single-molecule data generated by the dual-pore passes our aggressive quality filters for consensus generation and alignment. Pathologies for non-passing include too few tags or too sparse a tag pattern for unambiguous alignment. We expect that improvements in DNA sample preparation with increasing tag density will lead to large increases in this metric. Data throughput can be addressed with a dual-pore arrayed device with commensurate low-noise and multi-channel application-specific integrated circuits (ASICs). Finally, while our “zoom out” control logic allows for interrogation of *a priori* unknown tag numbers and tag patterns within molecules of *a priori* unknown length, we observed that the controller sometimes does not fully explore the molecule’s length. While this was not the case for *λ*-DNA with the 15-tag experiments, longer molecules may preferentially only be scanned near the end, and a failure to recapture the molecule may mean losing the opportunity to explore the entire tag set. Molecular dynamics simulations of molecule translocation and capture processes in two pore geometries may assist in further optimizing our scanning protocols to increase read-length, coverage and molecule capture efficiency.^32,39^

We demonstrated resolution of features separated by 150 bp at construct level and 166 bp vis-a-vis the *E. coli* genome reference map. This result is comparable to the 141 bp separation reported by Chen *et al* using a 5 nm diameter pore^3^ and superior to optical approaches which are limited to 1000 bp.^1^ We observed an average of 58.6% return to baseline between the detected peaks, therefore it is reasonable to expect that with sharper peak resolution from higher bandwidth recording the true resolution is less than 150 bp. We expect that optimization of the structure of the ODN tags and reduced DNA speed by tuning the competing voltages during flossing can also improve the spatial resolution of the dual-pore method.

In conclusion, we view our dual-pore approach as a first step towards genome scaling of solid-state nanopore technology. While genome mapping is a potential application, in the future we envision that the molecular feature used for barcoding (in our case ODN tags) can provide a scaffold for organizing, relative to the genome, the location of additional molecular motifs that might be discriminated based on size differences, such as nucleosomes or regulatory proteins, to provide a functional annotation/overlay on top of the sequence motif map. Labeling methods to differentially tag epigenetic sites can also be explored, e.g., across CpG islands to assay methylation. Somatic structural variations appear to have a major role in shaping the cancer DNA methylome.^40^ A tool that can capture genome-wide genomic and epigenetic alterations on single molecules, including changes to chromatin accessibility and methylation, would benefit research in cancers, aging-related diseases, and other conditions driven by such alterations.

## Methods

### Conjugation of oligodeoxynucleotide tags

We use two DNA substrates in this study, *λ*-DNA (New England Biolabs) and *E. coli* (K12 strain). Both are treated with nicking endonucleases as the first step in installing oligodeoxynucleotide (OdN) tags for dual-pore detection. The *λ*-DNA reagents are prepared starting with 2 *µ*g of commercially prepared DNA incubated with 25 units of Nt.BbvCI and Nb.BssSI to a final volume of 100 *µ*L in 1X 3.1 buffer (New England Biolabs). Nicking of genomic *E. coli* DNA was performed identically using Nb.BsrDI and 1X CutSmart (New England Biolabs) substituted for the two enzymes used to prepare *λ*-DNA. In both cases, the nicking reaction is incubated at 37°C for one hour. Nick translation was initiated by the addition of 5 *µ*L of 10 *µ*M dUTP-azide, dATP, dGTP, and dCTP, (ThermoFischer Scientific) and Taq polymerase (New England Biolabs) in 1X Standard Taq buffer. The mixture was incubated at 68° C for one hour at which point 3 *µ*L of 0.5 *µ*M EDTA was added to quench the reaction. The DNA was then purified by phenol/chloroform extraction and overnight ethanol precipitation. The dries pellet was washed with 70% ethanol, air dried, and resuspended in 500 mM NaCl/12.5 mM PO_4_ at pH 6. Synthetic OdNs with dibenzocyclooctynes (DBCO) moieties were added to the resuspended DNA to a final concentration of 1 *µ*M and reacted at 60°C overnight.^41^ Final reaction mixture was used in nanopore experiments directly following dilution in 2M LiCl to 1 ng/*µ*L. The OdN sequences can be found in Supplemental File 2. Both sequences had a 5^t^ DBCO moiety.

### Construction of 150 base pair tag-pair DNA

We generated a *λ*-DNA reagent with only 2 tags separated by 150 bp to test the resolution of the dual-pore instrument. We prepared the Cas9D10A nickase ribonucleoprotein (RNP) by incubating 1 *µ*L of 100 *µ*M annealed guide RNA and tracer DNA with 1*µ* L of 10 *µ*M Cas9D10A (IDT) in NEB 3.1 buffer for 10 minutes at RT then put on ice until use. The guide RNA sequences were CATTTTTTTTCGTGAGCAAT and AATTCAGGATAATGT-GCAAT for the 5^′^ and 3^′^ cut sites, respectively (Supplemental File 2). Both Cas9 RNPs were combined with the *λ*-DNA template to a final concentration of 100 nM RNP (each) and 1.56 nM *λ*-DNA and incubated at 37°C for 1 hour. The reaction was purified by phenol/chloroform extraction followed by ethanol precipitation and resuspended in deionized water. We then installed 60 nt tags at the nick sites using the methods described in **??**.

### Isolation of genomic *E. coli* DNA

The *E. coli* cells were grown in LB media overnight to stationary phase. The cells were harvested by centrifugation and genomic DNA was isolated using the Circulomics Nanobind HMW DNA extraction kit as per manufacturer instructions.

### Nanopore measurement

We performed nanopore experiments as previously described by our group.^18^ Briefly, the dual-pore chip was assembled in a custom fabricated flow cell. Our optimized flow cell uses 7 *µ*L of 1 ng/*µ*L of substrate DNA for each experiment. Ag/AgCl electrodes are fabricated to be used with the flow cell on a thin PET sheet that is positioned adjacent to the dual-pore chip and with electrodes in the relevant fluidic flow paths. The current and voltage signal was collected by Molecular Device Multi-Clamp 700B, and was digitized by Axon Digidata 1550. The signal is sampled at 250 kHz and filtered at 10 kHz. The tag-sensing and voltage control module was built on National Instruments Field Programmable Gate Array (FPGA) PCIe-7851R and control logic was developed and run on the FPGA through LabView.

### Tag Calling

The installed DNA tags cause a characteristic “spike” attenuation in the ionic current as the DNA moves through the nanopore sensors (Figure 2(b)). In online analysis, the flossing with zoom out logic requires detecting and counting tags. Detection is performed only in pore 2 current and when a tag creates a 70 pA or larger deviation in the 10 kHz signal compared to a moving average (2500 samples) filtered signal that emulates the DNA tag-free baseline signal. Tags that produce a deviation less than 70 pA are missed. In off-line analysis, to perform alignment, we need to determine where these tags translocate through the two nanopores. The process to determine the location of tags is the same for the ionic current from pore 1 and pore 2 and follows a 5 step process: (1) The current is inverted (making the downward spikes into upward ones); (2) An exponential fit is performed to detrend the transient in the baseline of the signalan (and this fit is subtracted from the signal); (3) the signal is mean-shifted and standardized; (4) a Gaussian filter is applied to the signal resulting in a smoothed representation and (5) a peak calling algorithm from the scipy library^42^ is applied.

### Data processing and filtering

We filtered the data for downstream use. This was done at the scan level, meaning a single molecule capture may have multiple scans (L-R or R-L recordings of the molecule) which pass or fail the filtering criteria. The process involves three steps, tag-count filtering, scan-length filtering and pore 1/pore 2 tag count imbalance filtering.

### Tag count filtering

For a molecule to progress to flossing, it must enter tug-of-war state, automatically filtering out small DNA molecules and material that is not linear dsDNA (for example DNA with a double strand break). However, occasionally DNA molecules or other material comes in contact with the dual-pore sensor during flossing that result in high noise signal. The tag detector will over-call the number of tags in these scans (often times detecting 100’s of tags erroneously), so we filter out scans with greater than 20 detected tags. For *E. coli* we remove scans without at least 4 tags, and for *λ*-DNA we remove scans with fewer than 3 tags.

### Scan length filtering

Short scans are typically false flossing triggers and are symptomatic of noisy or poor quality data. Similarly, extremely long scans are often due to stalling, molecule stiction in the pore or similar pathologies. We removed scans which are shorter than 1.6 ms and longer than 400 ms.

### Tag count imbalance filtering

We rely on detecting tags in both pore 1 and pore 2 to estimate velocity (Estimating scan velocity and linear distance between adjacent tags). We removed scans where the number of tags detected in pore 1 (*n*_1_) appreciably differed from the number of tags detected in pore 2 (*n*_2_) by calculating the relative difference in their counts and requiring:

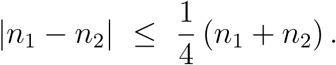

### Estimating scan velocity and linear distance between adjacent tags

The common chamber is the volume adjacent to both nanopores. We refer to “entering” tags as ones which are coming from the channel into the common chamber and “exiting” tags as ones leaving the common chamber into the channel. In the L-R direction the entering tags are detected in pore 1 and the exiting tags are detected in pore 2, whereas in the R-L direction the entering tags are detected in pore 2 and the exiting tags are detected in pore 1. Agnostic to scan direction, a tag is assigned a time-of-flight velocity *V*_TOF_ by dividing the tags entry-to-exit transit time into the known distance between the pores. This produces units of linear distance (nanometers) divided by time (microseconds) for *V*_TOF_. In detail, for each scan we iterate over the entering tags and look for corresponding exiting tags which would produce a *V*_TOF_ within the interval (0.4, 3.5) nm/*µ*s given the known pore-to-pore distance. Given multiple candidates, the first one in logical order is taken. Once a tag is paired, it is removed from consideration. For the purpose of generating a scan velocity profile, the time value assigned to each tags *V*_TOF_ value is at the point of that tags exit time. The scan velocity profile is the piece-wise linear curve that linearly interpolates between the computed *V*_TOF_ values, and is constant and equal to the first and last *V*_TOF_ values before and after they occur, respectively. Lastly, with either pore current, the linear distance between between two sequentially detected tags is computed by integrating the scan velocity profile over the inter-tag time period.

### Description of alignment algorithm

Individual scans are aligned to a reference map using an adapted version of the Smith-Waterman local alignment algorithm.^35^ There are two adaptations made for dual-pore distance data. First, instead of a substitution matrix, the score model described in Equation is used when determining the match score for a pair of tags and a length of nucleotides in the reference. Secondly, we must allow for incomplete tagging of the DNA molecule as well as spuriously detected tags. At a given step in the dynamic programming we search for the best match among interval covered up to the current position as well as the intervals covered using upstream tags as the start to the interval. More formally, let *s*_[*i*−*a,i*],[*j*−*b,j*]_ be the score *s*(*D*_*i*−*a,i*_, *G*_*j*−*b,j*_), where *D*_*i*−*a,i*_ is the estimated linear distance between tags indexed at *i* – *a* and *i, G*_*j*−*b,j*_ is the genomic interval distance between nicking sites indexed at *j* − *b* and *j*, and indices enumerate candidate restriction sites on the reference map. At each step in the dynamic programming recursion the match score is defined as:

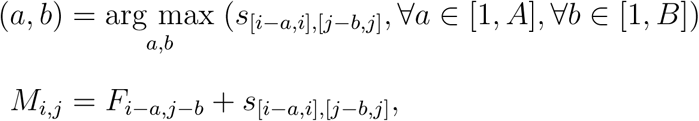

where *A* and *B* are nick-site counting parameters that define the index search size (default 3 for both), *F* is a dynamic programming matrix recursively defined below, and *M* is the optimized score. The recursions for “inserting” and “deleting” tags, as part of the dynamic programming search and which correspond to spuriously detecting tags or failing to detect a tag, respectively, use a cost based on the maximum score of the model. The maximum potential value for the score model is defined when the ratio 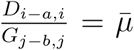, with 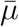 defined in Equation (1). Let *s*_max_ be the maximum score for the model. The recursion for inserting a tag is computed as

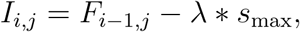

and the recursion for deleting a tag is similarly computed as

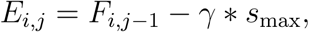

where *λ* and *γ* are parameters to the algorithm to adjust the relative cost of inserting a tag or deleting a tag. The default values are *λ* = 0.1, *γ* = 1.0. The full recursion for the dynamic programming is:

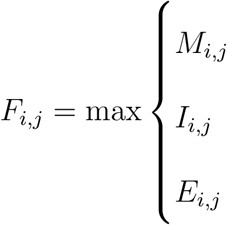

### Joining recapture alignments

Consensus alignments from a recaptured molecule may overlap (for example, Figure 6) or be disjoint. The DNA isolation procedure suggests that the fragments are longer than 50 kb (Figure S1). When the consensus alignments do not overlap we follow a procedure of grouping the consensus alignments from a single molecule when they have at least 2 scans of support and align to the reference within 250 kbp of each other. We then take the window with the highest scoring set of alignments as the final aggregated single molecule alignment. Our data contained 7 and 28 molecules with disjoint and overlapping consensus alignments, respectively. The mean distance between disjoint consensus alignments is 21.9 kb.

**Figure S1:**
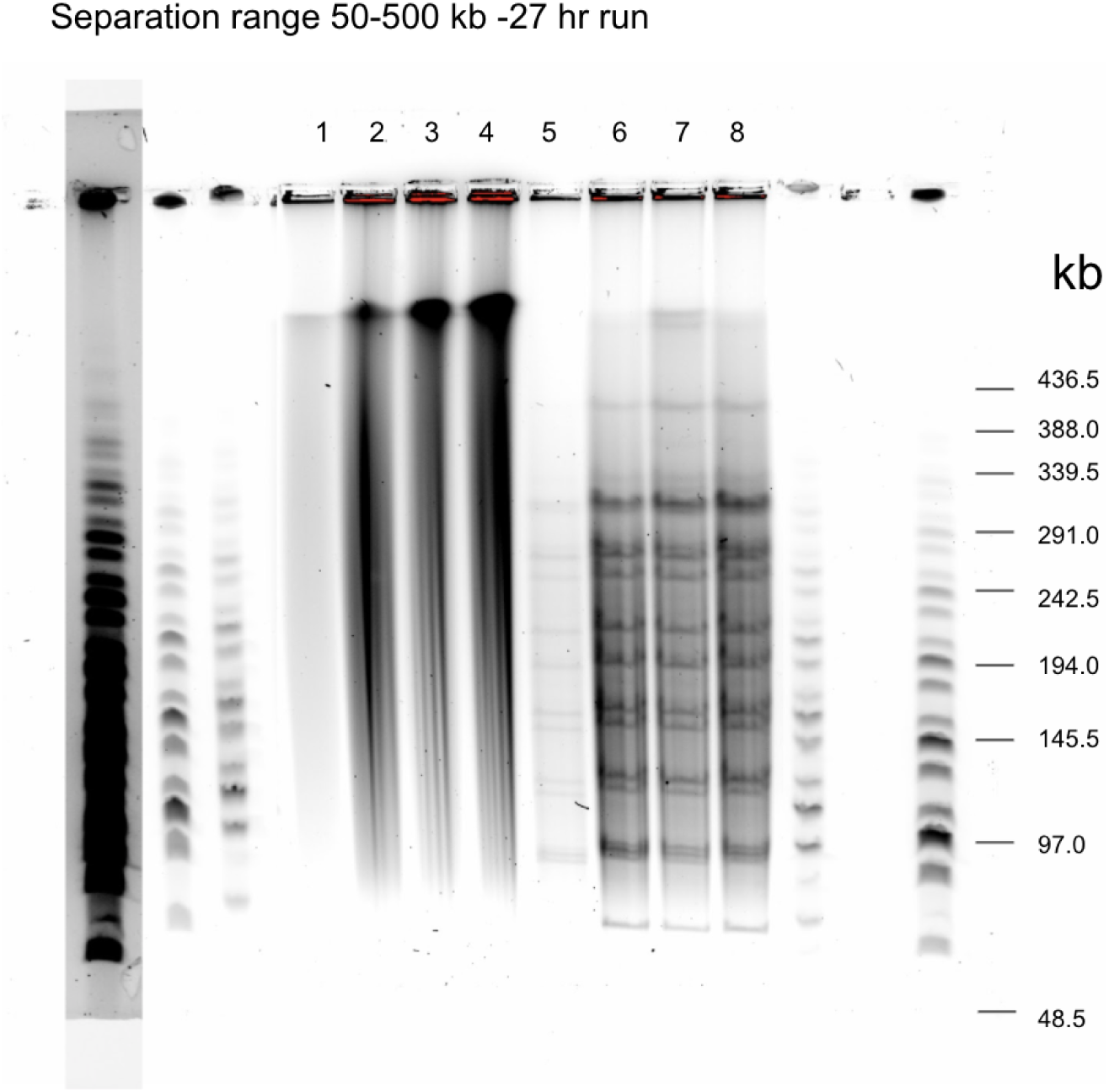
Pulse field gel of extracted *E. coli* genomic DNA showing high molecular weight input (lanes 1-4) and NotI digest of the same material (lanes 6-8).

### Calculating N50 of alignments

Let {𝒜_0_, 𝒜_1_, …, 𝒜_*n*_} be the set of all *n* + 1 consensus molecule alignments that met acceptance criteria. Let *𝓁* (𝒜_*j*_) be the net alignment length (in base pairs) defined as the maximum genomic coordinate minus the minimum genomic coordinate among the set of aligned tags, and computed for each consensus alignment 𝒜_*j*_, *j* = 0, …, *n* − 1. The net alignment length is thus a first-to-last tag distance, and does not include the length of the molecules outside of these tags. Next, let 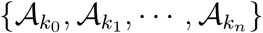 be the original set but now sorted in descending from largest net alignment length 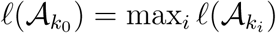 to the smallest net alignment length 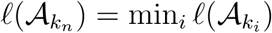. Define the cumulative sum of net alignment lengths as

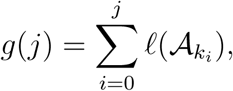

with the total alignment length denoted as *L* = *g*(*k*_*n*_). Note that the cumulative alignment length as a percentage of the total length of all alignments plotted in Figure 7(b) is *g*(*j*)*/L*. The 50% mark of the total alignment length is denoted *L*_50%_ = l0.5 × *L*J. Finally, we calculate the N50 as

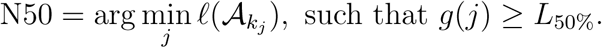

The N50 is the smallest net alignment length in the sorted list such that the cumulative sum up to that length is at least as big as the 50% mark of the total alignment length.

### Scan-space alignment

In order to estimate of the standard deviation for the DNA stretching factor we aligned scans from *λ* to a reference *scan*. For each molecule scan in the Phage Lambda dataset, we generate a synthetic reference scan by assuming the DNA stretching factor is equal to 0.34 nm/bp and the *V*_TOF_ for every tag is equal to the average for that scan.

The alignment procedure begins by taking one tag from each scan (called the anchor pair of tags, Figure S2, red line) and moving them into alignment, it then calculates time from the anchor tag *t*_*i*_ to every other distal tag *t*_*j*_ in the scan. The cost 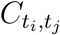 of aligning two tags is defined as the squared difference in their distance:

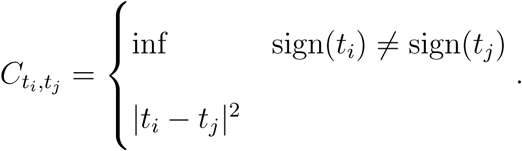

If 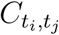 exceeds a user defined value (default 400*µ*s^2^) the tag is left unpaired. The distal tags are aligned (blue lines) using a dynamic programming algorithm analogous to global alignment^43^ to find the pairings which minimize the error. The first row and column of the dynamic programming matrix are initialized to the gap cost (a user-provided parameter) multiplied by the number of tags skipped on alignment. The recursion proceeds by taking the minimum of the difference in the distance between the two tags to their anchor, or incurring a skip cost added to the corresponding prior cell in the matrix. The minimum error over all pairwise combinations is taken from the bottom right cell of the matrix and a traceback is performed. Every pair of tags are attempted as the anchor pair. The alignment with minimum error over all anchor pairs is used as the final alignment. This procedure produces a mapping of scan intervals to genomic positions which we used to estimate the initial value of *σ*.

**Figure S2:**
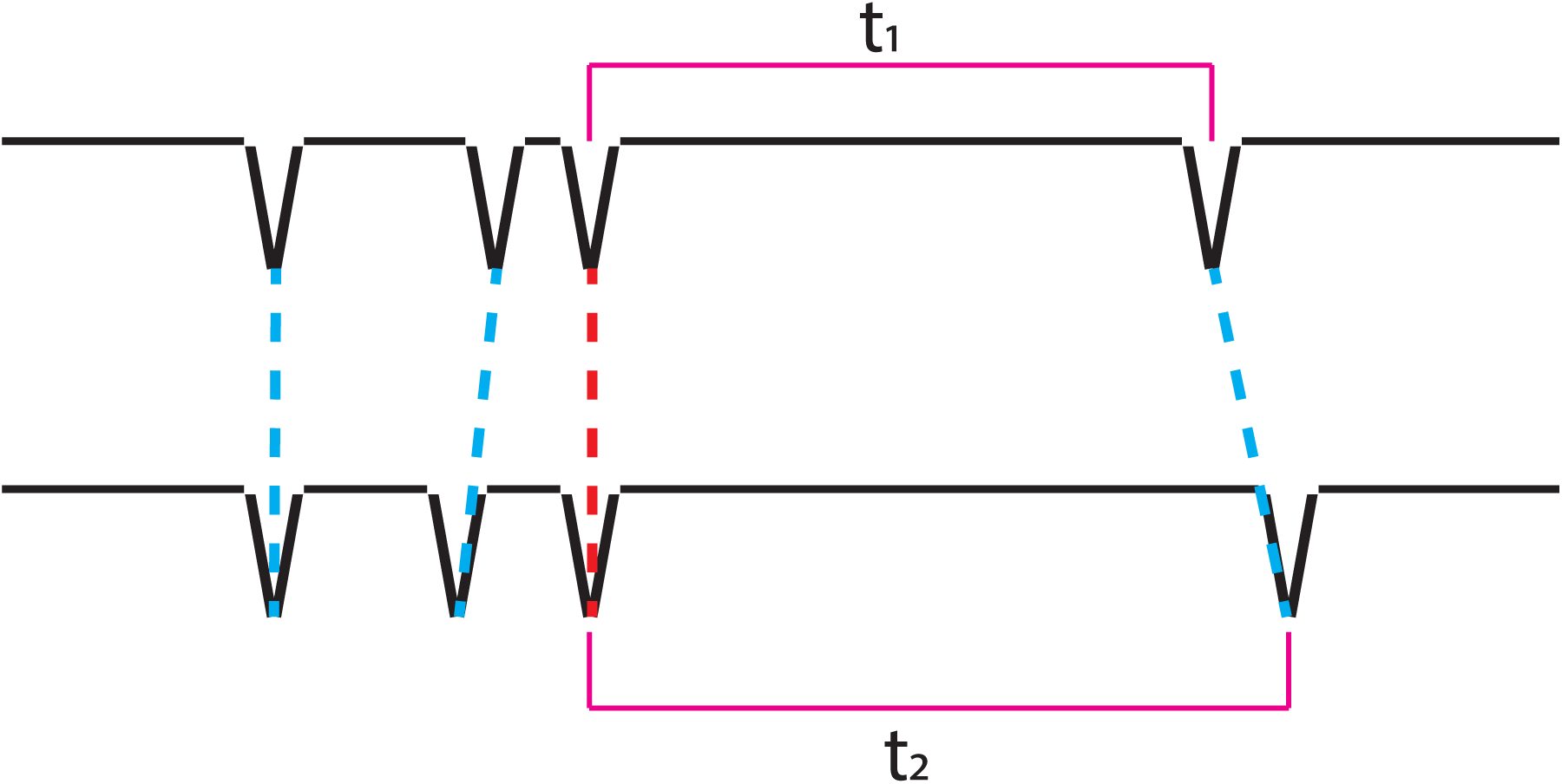
Cartoon of two scans being aligned in time space. Two tags are chosen as the anchor pair (red line) which are brought into alignment with each other. The remaining tags are paired (blue lines) using dynamic programming to minimize their error as defined by the difference in their (time-space) distance to the anchor.

