## Supplementary material for "Electronic Mapping of a Bacterial Genome with Dual Solid-State Nanopores and Active Single-Molecule Control": supplemental_files.tex

\section{Description of supplemental Files}

\subsection{Tandem Repeat Finder Output (Supplementary File 1)}

This archived folder contains the output from running Tandem Repeats Finder (version 4.09) on the \textit{E.
coli} genome.  The reference sequence used is included as well as the output containing the parameters for
running the program.

\subsection{Tag and guide RNA sequences}

This FASTA-formatted sequence file contains the nucleotide sequences of the 60nt tags used in this work. The
guide RNA sequences were used with the Cas9D10A nickase to install tags separated by 150\,bp on $\lambda$ DNA.
